# Abundance of Oligoflexales bacteria is associated with algal symbiont density independent of thermal stress in Aiptasia anemones

**DOI:** 10.1101/2023.04.14.536969

**Authors:** Emily G. Aguirre, Marissa J. Fine, Carly D. Kenkel

## Abstract

Many multicellular organisms, such as humans, plants, and invertebrates, depend on symbioses with microbes for metabolic cooperation and exchange. Reef-building corals, an ecologically important order of invertebrates, are particularly vulnerable to environmental stress in part because of their nutritive symbiosis with dinoflagellate algae, and yet also benefit from these and other microbial associations. While coral microbiomes remain difficult to study because of their complexity, the anemone Aiptasia is emerging as a simplified model. Research has demonstrated co-occurrences between microbiome composition and the abundance and type of algal symbionts in cnidarians. However, whether these patterns are the result of general stress-induced shifts or depletions of algal-associated bacteria remains unclear. Our study aimed to distinguish the effect of changes in symbiont density and thermal stress on the microbiome of symbiotic Aiptasia strain CC7 by comparing them with aposymbiotic anemones, depleted of their native symbiont, *Symbiodinium linucheae*. Our analysis indicated that overall, thermal stress had the greatest impact on disrupting the microbiome. We found that three bacterial classes made up most of the relative abundance (60-85 %) in all samples, but the rare microbiome fluctuated between symbiotic states and following thermal stress. We also observed that *S. linucheae* density correlated with abundance of Oligoflexales, suggesting these bacteria may be primary symbionts of the dinoflagellate algae. The findings of this study help expand knowledge on prospective multipartite symbioses in the cnidarian holobiont and how they respond to environmental disturbance.

## INTRODUCTION

Biological organisms, from single cells to ecosystems, are influenced by symbiotic interactions (Boucher, 1985; University of Massachusetts Amherst Massachusetts Lynn Margulis, Margulis and Fester, 1991; Smith and Szathmary, 1997; Sachs *et al*., 2004). Although symbioses are generally modeled and studied as two-way interactions, multipartite symbioses are also ubiquitous and have been well-documented in plants (Miransari, 2011; Antunes and Goss, 2015; Adeniji, Babalola and Loots, 2020; Afkhami *et al*., 2020), humans (Wahida, Tang and Barr, 2021), and invertebrates (Worthen, Gode and Graf, 2006; Ffrench-Constant, Eleftherianos and Reynolds, 2007; Chaston and Goodrich-Blair, 2010; Stock, 2019). These multi-partner interactions can provide the host with essential metabolites, amino acids, and vitamins (Cleveland, Alan Verde and Lee, 2011; Adeniji, Babalola and Loots, 2020). Additionally, they can also confer defense (Zan *et al*., 2019), facilitate host morphogenesis (Wichard, 2015) and aid resilience under environmental stress (Gupta *et al*., 2021; Santoyo *et al*., 2022).

Understanding the impact of multi-partner associations on the health and survival of marine invertebrates is crucial given they are among the most susceptible animals to the impacts of climate change (Mather, 2013; Lam *et al*., 2020). The global decline of corals, which are cnidarian hosts that harbor intracellular populations of dinoflagellate algae in the family Symbiodiniaceae and other microbial associates (Pandolfi *et al*., 2003) is already underway and is predicted to worsen with climate change (Hoegh-Guldberg *et al*., 2007; Allemand and Osborn, 2019; Kleypas and Kleypas, 2019). Bleaching, which involves the expulsion of symbiotic algae in cnidarians resulting in the loss of color, can be induced by several factors, although elevated temperature, as noted by (Douglas, 2003)), is the most common cause. Yet there is also abundant variation in coral thermal tolerance evidenced by differences in bleaching among species, populations, and individuals (Dixon *et al*., 2015; Thomas *et al*., 2018; Drury, 2020). Collectively, these observations have led to the "Coral Probiotic Hypothesis” (Reshef *et al*., 2006) which is based on the notion that symbiotic relationships with bacteria can increase coral resilience (Peixoto *et al*., 2017).

The cnidarian-dinoflagellate symbiosis is better characterized in the literature than potential cnidarian-bacterial symbioses, and it remains unclear whether environmental or internal host factors, like host genetics or microalgal symbiont type, modulate the composition of cnidarian microbiomes (Bourne, Morrow and Webster, 2016; van Oppen and Blackall, 2019; Barno *et al*., 2021). One understudied theory postulates that well-known cnidarian-bacterial associates may actually be primary associates of Symbiodiniaceae, in both free-living and in-hospite states (Ritchie, 2012; Bernasconi *et al*., 2019; Matthews *et al*., 2020). Global datasets suggest that the identity of Symbiodiniaceae may contribute to structuring coral and anemone microbiomes (Bernasconi *et al*., 2019). For instance, susceptibility to *Vibrio* pathogens was higher in *Acropora cytherea* corals hosting *Symbiodinium* than *Durusdinium* (formerly Clade A and D Symbiodinium, respectively) (Rouzé *et al*., 2016). Additionally, the abundance of diazotrophs in *Montipora* corals also correlated with algal symbiont type (Olson *et al*., 2009).

Similarly, unique microbiomes were identified in symbiotic and aposymbiotic Aiptasia anemones (Herrera *et al*., 2017) indicating presence of the symbiont influences the microbiome. However, insufficient evidence exists to verify this hypothesis and tracking bacteria in adult corals is nearly impossible, due to their high bacterial diversity (Blackall, Wilson and van Oppen, 2015), which poses a challenge for distinguishing obligatory and facultative bacterial symbionts. Furthermore, selectively eliminating holobiont members empirically is not feasible since reef-building corals cannot survive without their algal symbionts, who provide vital sugars and nutrients (Weis, 2008). The anemone Aiptasia (*Exaiptasia pallida*, *sensu stricto*) is a tractable model for studying the cnidarian microbiome as it harbors fewer bacterial OTUs, with diversity estimated to be around 1-2 orders of magnitude lower than their coral relatives, (Röthig *et al*., 2016; Herrera *et al*., 2017). Aiptasia are easy to maintain, engage in a nutritive symbiosis with Symbiodiniaceae similar to corals, and can reproduce asexually (Baumgarten *et al*., 2015; Weis, 2019). Unlike their coral relatives, Aiptasia can be rendered aposymbiotic (free of their dinoflagellate algae) in laboratory conditions and lack a calcitic skeleton, which facilitates experimental manipulation (Lehnert *et al*., 2014).

While associations between microbial community composition and the abundance and/or diversity of algal endosymbionts is consistent with the hypothesis that some microbes are primary associates of Symbiodiniaceae, these patterns cannot be distinguished from passive commensal relationships. Additionally, general stress may be responsible for changes in the host-photosymbiont relationship, leading to alterations in metabolite production and resulting in different selective pressures that favor distinct groups of commensals. For example, Symbiodiniaceae taxa are known to differ in their metabolite production (Camp *et al*., 2022). *Durusdinium* translocates less carbon to hosts than *Cladocopium (*Cantin *et al., 2009)* under thermal stress but *Cladocopium* translocates more carbon and nitrogen to hosts during non-stressful conditions (Pernice *et al*., 2015), which could impact the consortium of microbiota in the holobiont. The host-photosymbiont relationship can also be affected by mild bleaching, which is another general stress response (Ortiz, Gomez-Cabrera and Hoegh-Guldberg, 2009). Here, we aimed to distinguish the effects of the stress response and that of symbiont density on microbial communities while controlling for host and symbiont genetic diversity. To achieve this, we used the emerging model organism Aiptasia clonal strain CC7, which harbors *Symbiodinium linucheae (*Starzak *et al.,* 2014; Baumgarten *et al.,* 2015) and conducted a comparison between the microbiomes of symbiotic anemones undergoing mild bleaching and aposymbiotic anemones, which were presumed to lack the microbes typically associated with *Symbiodinium*.

## MATERIALS AND METHODS

### 2.1 Aiptasia rearing

Aiptasia anemones, clone strain CC7 (obtained from Dr. Cory Krediet, Eckerd College, FL, USA) were used in this study. Anemones were kept in 0.5 L polycarbonate tanks, filled with 0.2 μm filtered seawater (FSW) from Catalina Island (Catalina Water Co., Long Beach CA, USA) and maintained at 25°C on a light/dark (14:10 h) cycle under 12-20 µmol photons m -2 s -1. Animals were maintained in these common garden conditions at USC with weekly feeding (frozen brine shrimp, *Artemia salina*) and water changes for 24 months. A subset of Aiptasia animals were rendered aposymbiotic by menthol-induced bleaching (Matthews *et al*., 2016) four months prior to experimental trials. Briefly, a 1.28M menthol solution was prepared (20% w/v in ethanol) and added to polycarbonate tanks containing FSW for a final concentration of 0.19 mmol l -1. Anemones were placed in the menthol solution for 8 h during the light period of the 14:10, light: dark cycle. The animals were then transferred overnight to tanks containing a final concentration of 0.10 M DCMU (3-(3,4-dichlorophenyl)-1,1-dimethylurea), an algicide and photosynthesis inhibitor. This was repeated for four consecutive days. At the end of the fourth day, anemones were placed in black-out tanks with no light exposure and allowed to recover for three days with one feeding. After the three-day recovery, the process was repeated once more and bleaching status was confirmed using a fluorescent microscope. An absence of red autofluorescence from dinoflagellate chloroplasts was noted and these aposymbiotic anemones were transferred to black-out tanks filled with 0.2 µm FSW and maintained in the dark for three months with the same feeding and water change regime as the symbiotic anemone stock.

### 2.2 Experimental design and sampling

Symbiotic and aposymbiotic anemone of approximately 2.5 cm length were selected for the 6-day thermal stress experiment. Aposymbiotic Aiptasia were added to the experimental design to act as a control for distinguishing baseline host stress responses from cnidarian-algal microbiome shifts. These aposymbiotic anemones were acclimated to the same light/dark (14:10 h) cycle as symbiotic anemones one week prior to the experiment start date. In addition, all anemones were starved for 2 weeks prior to sampling to avoid prey (shrimp, *Artemia sp.*) contamination.

Six 0.5 L tanks containing six symbiotic Aiptasia (n=36) and four 0.5 L tanks containing six aposymbiotic Aiptasia (n=24) were distributed evenly between experimental conditions (25 °C vs 32 °C, Fig. 1). The treatment temperature was set to 32 °C due to prior observation of slight bleaching and expulsion of *Symbiodinium* in Aiptasia at this temperature (Perez, Cook and Brooks, 2001)). To maintain and manipulate temperature, tanks used in the heat stress treatment were placed in 10 L bins containing two SL381 submersible water pumps, two 100W aquarium heaters, a HOBO temperature logger, and a digital, waterproof thermometer. Acclimatization of Aiptasia was reached by exposing them to a gradual temperature ramp over the course of four days. Temperature was increased from 25 ± 0.5 °C to 27 ±0.5 °C the first day, to 30 ± 0.5 °C the second day, 32 ± 0.5 °C the third day and 33 ± 0.5 °C on the fourth day. Once Aiptasia were acclimated to the elevated temperature treatment, treatment was maintained at an average of 32 ± 0.5°C for 6 days with FSW changes every 2 days, in both control and heat-stress tanks. At the end of the 6-day exposure period, each anemone was rinsed three times with FSW in individual 60 x 15 mm petri dishes, placed in a 1.5 mL microcentrifuge tube and frozen at -80 °C, until processing.

**Figure 1.**
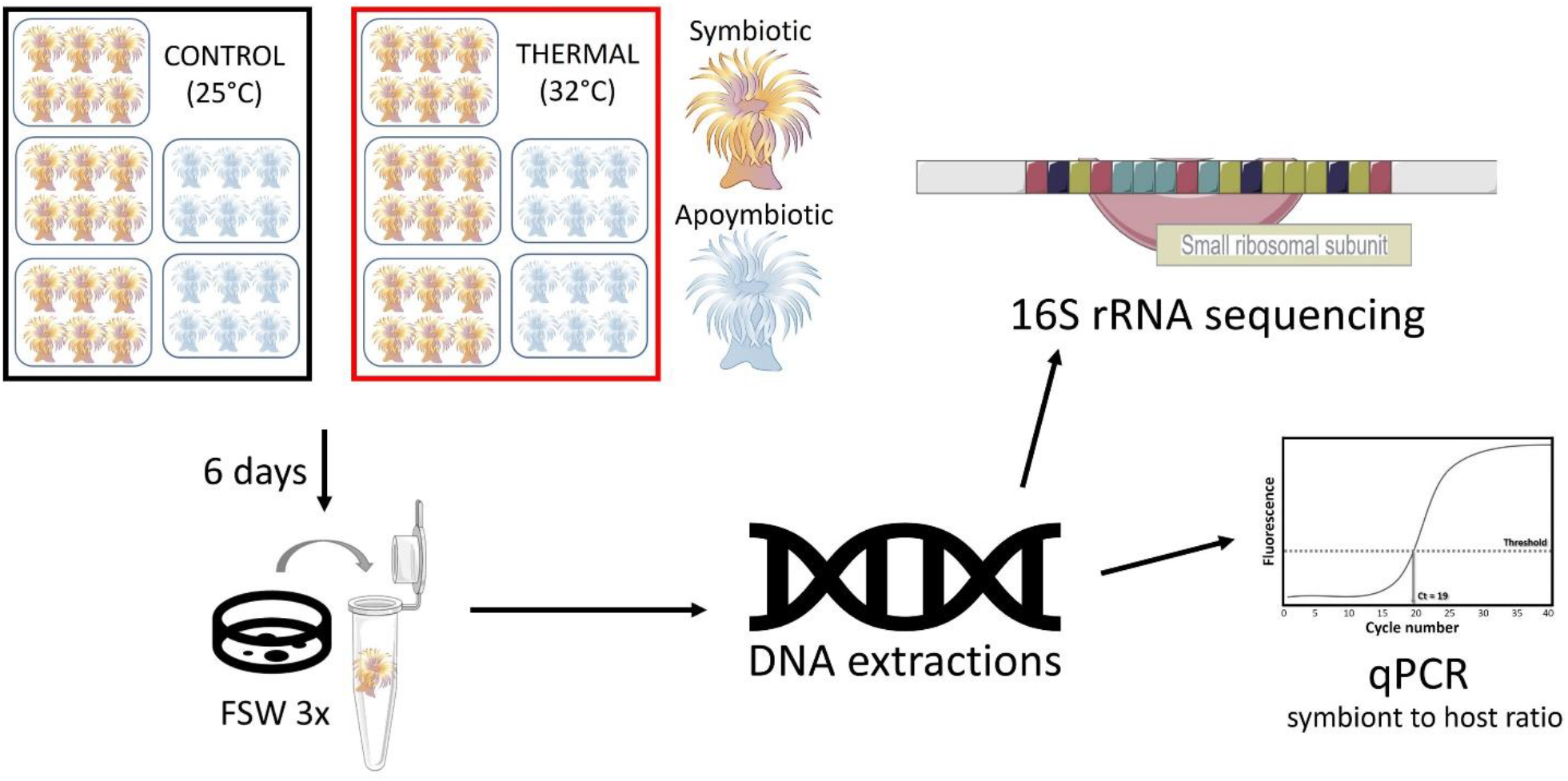
Study design for the mild thermal stress experiment. Each treatment condition received 5 tanks (3 tanks x 6 symbiotic anemones and 2 tanks x 6 aposymbiotic anemones; n= 30 per treatment). Following a 7-day ramping period, the peak temperature exposure continued for 6 days, and sampling was conducted on the last day. Each anemone was rinsed with filtered seawater (FSW) thrice, deposited into a microcentrifuge tube and stored at -80 °C until processing. DNA extractions were performed followed by 16S rRNA amplicon sequencing. Samples with remaining DNA were used in qPCR assays to determine symbiont *(Symbiodimitm)* to host (Aiptasia) ratios.

### 2.3 DNA extractions and 16S rDNA sequencing

Individual anemone DNA was extracted by ethanol precipitation as detailed previously (https://openwetware.org/wiki/Ethanol_precipitation_of_nucleic_acids) with some modifications. Briefly, animals were placed in sterile, 2 mL polypropylene screw cap tubes (Merck KGaA, Germany) containing a thin layer of Zi/Si beads (100-500 mm diameter), 200 μL lysis buffer AP1 from a DNEasy Power Plant Kit (Qiagen, Germany), 2 μL RNAse (100 mg/mL, stock), and 2 μL Proteinase K (20 mg/mL, stock) and incubated at 55 °C for 10 min in a temperature-controlled water bath. Samples were then homogenized using the Omni bead beater (Omni International, USA) at 6.3 m/s, for 2 cycles of 30 seconds. The samples were then centrifuged for 5 min at 14,000 rpm and the proteinase K enzyme was heat-inactivated on a heat block (78-82 °C) for 3 min. The supernatant was transferred to a 1.5 mL microcentrifuge tube.

Ethanol precipitation was performed as described (https://openwetware.org/wiki/Ethanol_precipitation_of_nucleic_acids) and the pellet was resuspended in 40 μL elution buffer (GenElute Bacterial Genomic DNA Kit, Merck KGaA, Germany). DNA was purified using the Zymo DNA Clean & Concentrator Kit (Zymo Research, USA) following the manufacturer’s instructions.

Amplification of the V5/V6 region of the 16S rRNA gene was done using the 784F and 1061R primer set, which amplifies approximately 277 bp fragments with minor cross-amplification of host mitochondria and microalgal symbiont chloroplasts (Andersson *et al*., 2008; Bayer *et al*., 2013). Briefly, 50 ng of DNA were amplified in 25 ul reactions using 0.25 μL of 2,000 units/mL Q5 High-Fidelity DNA Polymerase (New England BioLabs Inc., Germany), 0.5 μL of 10 mM each dNTP, 0.25 μL of 10 μM forward and reverse primer each, 0.25 μL BSA 100X (New England BioLabs Inc., Germany), 5.0 μL 5X Q5 PCR Buffer (New England BioLabs Inc., Germany), 13.5 μL nuclease-free water, and 5 μL of 10 ng/μL DNA. PCR conditions were as previously described in Bayer et al. (2013) with the following modifications: 26 cycles of denaturation at 95°C for 60 sec and annealing at 55°C for 60 sec. 5 mL of the 16S rRNA amplified sample was added to a second round of 3-step PCR to incorporate sample-specific Illumina barcodes using an amplification profile of: 98°C 0:30, (98°C 0:10 | 59°C 0:30 | 72°C 0:30) x 4 cycles, 72°C 2:00. The barcoded samples were pooled in equimolar amounts (1 ng/μL) and sent for 250-bp paired-end sequencing on Illumina’s MiSeq v2 PE 250 platform (USC Norris Comprehensive Cancer Center Molecular Genomics Core (NCCC), USA).

### 2.4 16S rDNA sequencing and bioinformatic analysis

We successfully extracted DNA and sequenced the 16S rRNA gene of 59 individual anemones (initial n=60) but seven samples were discarded from the dataset due to low read yields (< 2,000). Paired-end reads were demultiplexed by the sequencing facility (USC NCCC Molecular Genomics Core) and quality checked with FastQC (Andrews, 2010). An amplicon sequencing variant (ASV) table was was generated with DADA2 (Callahan *et al*., 2016) in R (R Core Team 2020) using the default filtering parameters (truncLen=c (240,160), maxN=0, maxEE=c(2,2), truncQ=2, rm.phix=TRUE, compress=TRUE, multithread=FALSE). Taxonomy was assigned using the Ribosomal Database Project Classifier (Wang *et al*., 2007) along with the SILVA SSU version 138.1 database, formatted for DADA2 (DOI/10.5281/zenodo.4587955). Chloroplast and mitochondria sequences were removed using the R package, Phyloseq (McMurdie and Holmes, 2013). The dataset yielded uneven library sizes, so samples were rarefied to an even depth of 26, 481 reads (Gloor *et al*., 2017; Weiss *et al*., 2017). Following rarefaction, the dataset consisted of 1, 377, 012 reads (Table A1). All statistical analyses and visualizations were conducted in R (R Core Team, 2020). Taxa plots, alpha diversity (Chao1 index), and beta-diversity visualizations were conducted using the Phyloseq (McMurdie and Holmes, 2013), ggplot2 (Wilkinson, 2011), and Microbiome (Lahti and Shetty *et al*, 2019) packages. Differential tree matrices were generated and visualized using the Metacoder (Foster, Sharpton and Grünwald, 2017) package. Statistical output for differential heat trees can be found at Zenodo, <u>doi.org/10.5281/zenodo.7693398</u>.

Additional statistical analyses were conducted using the vegan (Dixon, 2003) package. A one-way ANOVA test was conducted on rarefied data, comparing alpha-diversity data (Chao1 scores) between the microbial assemblages of symbiotic and aposymbiotic anemone groups in different treatments (control vs thermal stress). Post-hoc pairwise comparisons were done using Tukey’s HSD. Beta-diversity was visualized using a PCoA plot employing the weighted-Unifrac metric. The adonis2 function in vegan was used to conduct pairwise Permutational Multivariate Analysis of Variance (PERMANOVA) of microbial assemblage dissimilarities between treatment groups. Adonis2 was used due to even homogeneity of variances in all pairwise comparisons tested.

As the SILVA SSU v138.1 database only assigned some bacterial taxonomy to Order, we conducted a separate phylogenetic analysis of ASVs classified in the order Oligoflexales. A Bioconda (Grüning *et al*., 2018) environment was used for this analysis. The filtered Phyloseq dataset was subsetted to include only Oligoflexales reads and resulting ASVs were then transferred to a fasta file and a standard NCBI nucleotide blast (Sayers *et al*., 2023) was performed on the web interface, optimizing for highly similar sequences (megablast). The ASVs exhibited high quality (Table A2) matches to three uncultured bacterium clones, originating from one microbial survey (Randle *et al*., 2020) conducted on Aiptasia strain CC7 (GenBank: MK571601.1/ MK571569.1) and Aiptasia strain H2 (GenBank: MK571216.1), as well as one uncultured proteobacterium clone (GenBank: FJ425635) found in the microbiome of scleractinian coral *Orbicella* (formerly *Montastrea*) *faveolata*. We queried our nine Oligoflexales ASVs and these previously published sequences with 16S rDNA sequences from 2 confirmed members of the Oligoflexales order (GenBank accession numbers AB540021.2 and OW948931.1) and ten members (7 families) of the Bdellovibrionota phylum. Sequence alignment was performed using the MUSCLE algorithm version 5.1 (Edgar, 2004) and a phylogenetic tree was constructed by maximum likelihood with ultrafast bootstrap (n= 1,000 replicates) in IQ-TREE version 2.2.0.3 (Kalyaanamoorthy *et al*., 2017; Minh *et al*., 2020). The resulting phylogenetic tree was visualized and annotated using the integrated web editor interface, Interactive Tree of Life (ITOL, https://itol.embl.de/).

### 2.5 Symbiont to host (S/H) cell ratio qPCR

To assess the effects of thermal stress on symbiont density in Aiptasia, we analyzed symbiont to host cell ratios (Mieog *et al*., 2009) using nuclear ribosomal protein L10 as a reference gene for the host, Aiptasia (Poole, Kitchen and Weis, 2016) and the actin locus as a target in *Symbiodinium* (Palacio-Castro 2019). Nuclear ribosomal protein L10 primers were previously validated for Aiptasia specificity in qPCR assays (Poole, Kitchen and Weis, 2016) and used as a reference gene due to stable expression in Aiptasia (Kitchen and Weis, 2017). Previous qPCR assays targeting the actin locus gene in *Symbiodinium* sp. (clade A) showed amplification specificity with an estimated copy number for the actin locus at ∼9 per cell (Palacio-Castro 2019). The primers used in this study for the host were 400nM nrp_L10-F (5’-ACGTTTCTGCCGTGGTGTCCC-3’) and 400 nM nrp_L10-R (5’-CGGGCAGCTTCAAGGGCTTCA-3’). Primers used for *Symbiodinium* symbionts were 300 nM Aact_F (5’-ATGAAGTGCGACGTGGACAT-3’) and 200nM Aact_R (5’-GGAGGACAGGATGGAGCCT-3’). All qPCR assays were performed on the Agilent AriaMx qPCR Machine (Agilent, USA). Each reaction totaled 20 μL volumes, using 10 μL Brilliant III Ultra-fast SYBR qPCR Master Mix (Agilent, USA), 6.1 μL Milli-Q water, 0.8 μL per primer (forward and reverse, final concentrations listed above), 0.3 μL 1:500 reference dye (SYBR) and 2 μL of template DNA (concentration range between 5-10 ng/ μL). The thermal profile was 50 °C 2:00, 95 °C 10:00 (95 °C 0:10 | 60 °C 1:00 | 72°C 0.20) x 40 cycles + melt curve profile of (95 °C 0:30 | 65 °C 0:30 | 95 °C 0:30) x 1 cycle. Due to low remaining DNA after 16S rRNA library preparation and sequencing, we used 15 aposymbiotic replicates and 21 symbiotic replicates (Table A3) for the qPCR assays. Each sample was assayed in duplicate, per target primer set. Cycle threshold (Ct) values were calculated by Agilent AriaMx qPCR machine when the first amplification cycle in a reaction exceeded the fluorescent baseline. All aposymbiotic anemones in the control (but not those in the thermally stressed samples) exhibited non-target amplification for the actin primer set, and upon analysis of the melt curve, we noticed a distinct peak between 83-84 °C for these samples, yet all other positive amplification reactions exhibited a distinct peak between 85-87 °C. We surmised cross-amplification of an Aiptasia locus in the absence of a *Symbiodinium* target. Sanger sequencing of the different products revealed that actin samples with a melt product between 85-86 °C exhibited high quality matches to the actin gene locus in *Symbiodinium* spp. (GenBank: AB231899.1, NCBI nr BLAST, megablast, e-value= 8e-86, bit score= 329) whereas no matches were identified for the lower melt product samples, likely due to a high abundance of Ns in the sequences. Therefore, actin reactions exhibiting melt products < 84 °C were assigned a cycle number of 40 to account for this non-specific amplification (Table A4). Ct values were averaged between technical replicates and symbiont to host ratios were calculated using the formula ((2^ (Ct host)/(Ct sym))∗2) based on host/symbiont target ploidy (Aiptasia, host = 2, *Symbiodinium*, symbiont = 1) (Cunning and Baker, 2013; Palacio-Castro 2019).

### 2.6 S/H ratio statistical analyses

A dataset containing only symbiotic Aiptasia with S/H ratios, their alpha-diversity scores (Shannon, Chao, Observed, Fisher and Simpson) and select bacterial abundance counts (Oligoflexales and Staphylococcus) was used to generate a Pearson’s correlation matrix using the corrplot package in R (Wei and Simko, 2021). The lm command was used for regression analysis of Oligoflexales abundance counts on S/H ratio in symbiotic anemones. We implemented a linear mixed effects model with a fixed effect of treatment and a random effect of tank to test whether S/H ratios were reduced in symbiotic anemones response to heat treatment using the nlme (Pinheiro and Bates, 2023; Bates and Pinheiro, 1998), and lme4 (Bates *et al*., 2015) packages.

## RESULTS

### 3.1 Microbial assemblages in Aiptasia

After ASV calling with DADA2, the sequence table yielded 4, 996, 356 reads and after chloroplast and mitochondria removal, 4, 984, 116 reads and 4, 471 ASVs remained (Table A1). Rarefaction and filtering of taxa occurring at least 3 times in more than 4 samples (the minimum number of replicates per tank) yielded a final dataset of 1, 338, 082 reads and 774 ASVs. The dataset was dominated by Alphaproteobacteria (66 %), Gammaproteobacteria (19%), and Bdellovibrionota (4%) (Fig. A1). All other taxonomic classes were present in abundances < 3%, except for Oligoflexia, which occurred at 4% relative abundance in symbiotic anemones under control conditions (Fig. A1). The predominant genera in all samples were *Cognatishimia*, an unnamed bacterium from the family Rhodobacteraceae and *Alcanivorax* (Fig. A2).

### 3.2 Community dynamics differ by treatment and symbiotic state

Alpha diversity was assessed by estimating species richness using the Chao1 index. Lower within-sample diversity was observed in symbiotic anemones exposed to heat, but within-sample diversity was consistent between other treatment groups (Fig. 2a). Although alpha diversity differed between groups on average (ANOVA, p=0.009, Table A5), significant differences were detected for only one pairwise comparison: heat-stressed symbiotic anemones and control symbiotic anemones (Tukey multiple comparison of means, p=0.016, Table A5). A marginal pairwise difference was detected between symbiotic and aposymbiotic anemones under heat stress (Tukey multiple comparison of means, p=0.05, Table A5).

**Figure 2.**
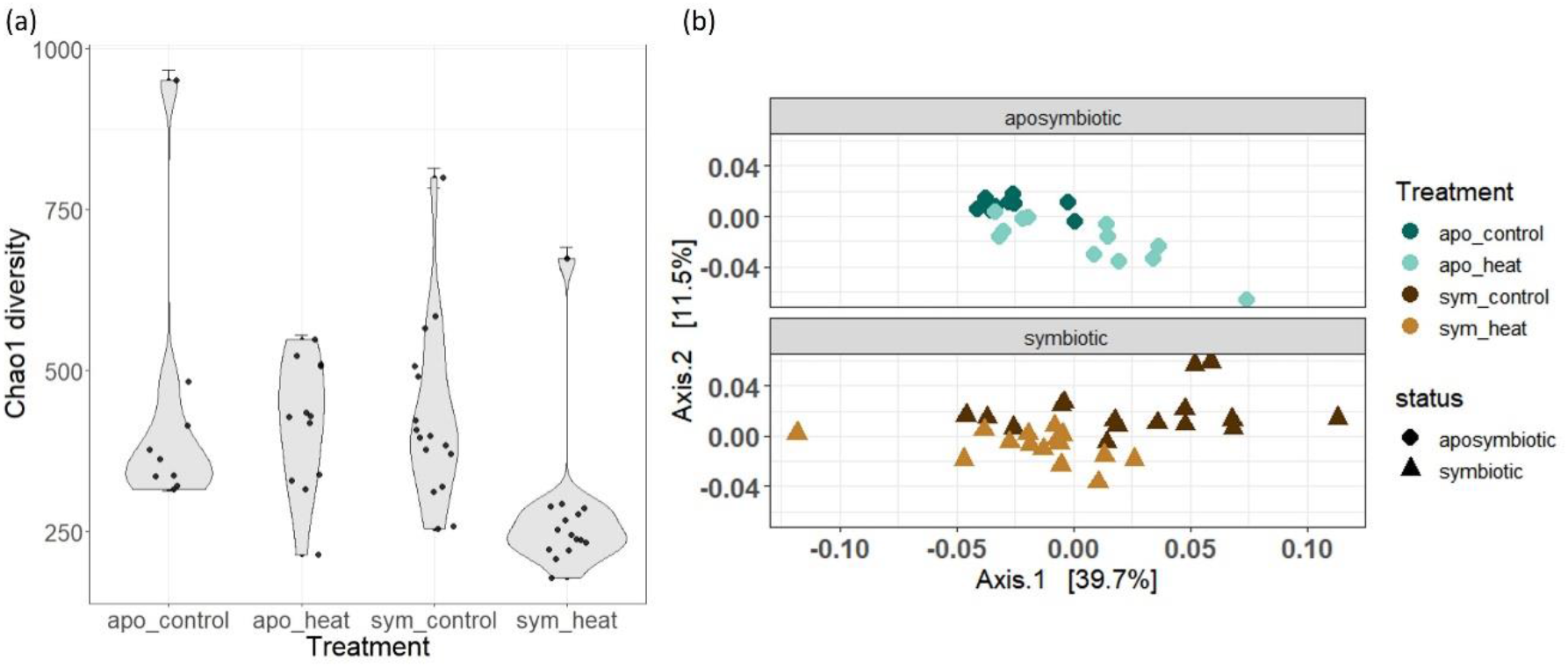
Alpha diversity and beta diversity differences of microbial assemblages in Aiplasia. (a) Alpha diversity index, Chaol, by treatment group. Standard error for Chaol is represented by the error bars, (b) Bela differences by weighted Unilrac, principal coordinates of analysis (PCoA) on aposymbiotic samples (control, dark blue circles vs heat stressed, light blue circles) and symbiotic (control, dark brown triangles vs heat stressed, light brown triangles).

A principal coordinate analysis (PCoA, weighted Unifrac) also revealed differences in beta diversity between symbiotic and aposymbiotic pairs in response to treatment (Fig. 2b). Namely, beta diversity in heat-stressed symbiotic anemones converged whereas beta diversity in control symbiotic anemones did not (PERMANOVA, p= 0.001, Table A6), indicating microbial community composition in symbiotic anemones became more similar to each other after heat treatment. The opposite pattern was observed in aposymbiotic anemones: convergence was observed in the control group and divergence in the heat treatment (Fig. 2b, PERMANOVA, p=0.001, Table A6). Differences in beta diversity were also observed between symbiotic and aposymbiotic animals under control conditions (ANOSIM R= 0.28, p = 0.003, Table A6).

We further explored taxa responsible for beta diversity disparities by generating a differential heat tree and visualizing statistically dissimilar taxa between pairwise comparisons. We detected differential abundance of several taxa (Wilcoxon Rank Sum tests, FDR-adjusted p-values < 0.05. Statistical output: Zenodo, <u>doi.org/10.5281/zenodo.7693398</u>) but most noticeably Firmicutes, Oligoflexales, Oceanospirillales, Planctomycetes and Alteromonadales (Fig. 3).

**Figure 3.**
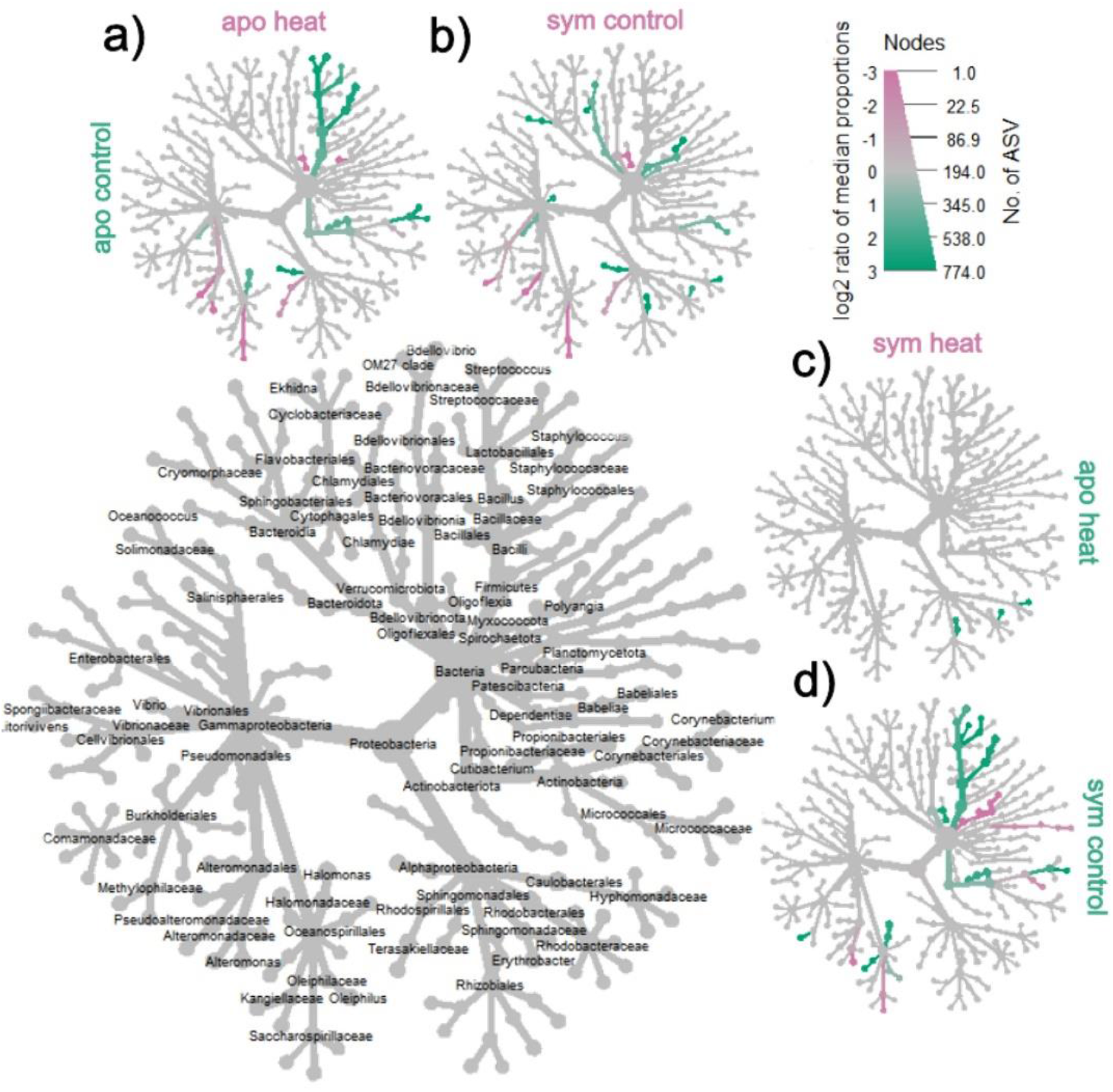
Differential heat trees illustrating pairwise comparisons between groups of interest. Amplicon data was used to visualize microbial taxonomic composition in Aiptasia, using the R package, Metacoder (Foster et al. 2017). The bigger tree with taxon labels on the lower left serves as a key for the smaller pairwise-comparison trees surrounding it a) aposymbiotic control vs aposymbiotic heat, b) aposymbiotic control vs symbiotic control, c) symbiotic heat-stressed vs aposymbiotic heat-stressed and d) symbiotic heat-stressed vs symbiotic control. Taxon color (diverging scheme from pink to green) is represented by log-2 ratio of median proportions of reads observed by treatment group. Significantly differentially abundant taxa, determined by Wilcoxon rank sum tests followed by FDR correction, colored in pink arc more prominent in the groups shown in the columns and those colored in green arc more prominent in the groups shown on the rows, e.g., Oligoflexales are significantly more abundant in symbiotic control (green) anemones than symbiotic heat-stressed but Myxococcota are enriched in symbiotic heat-stressed (pink). Size of tree nodes corresponds to ASV richness, as denoted in the color and size key in the upper right. Statistical output of differential abundance analysis is archived at Zenodo, doi.org/10.5281/zenodo.7693398

Firmicute abundances were higher in aposymbiotic and symbiotic controls (Fig 3a, 3d), and Oligoflexales abundances were highest in symbiotic anemones under control conditions (Fig. 3b). Bacteria from the family *Methylophilaceae* exhibited a similar pattern as Oligoflexales but overall abundance was low, and the difference between aposymbiotic and symbiotic controls was only ∼ 100 raw counts. In contrast, Oligoflexales abundances in the aposymbiotic and symbiotic controls differed by 1-3 orders of magnitude (Fig. 4b). Alteromonodales and Oceanospirillales abundance both decreased in heat-stressed anemones, regardless of symbiotic state (Fig. 3c).

**Figure 4.**
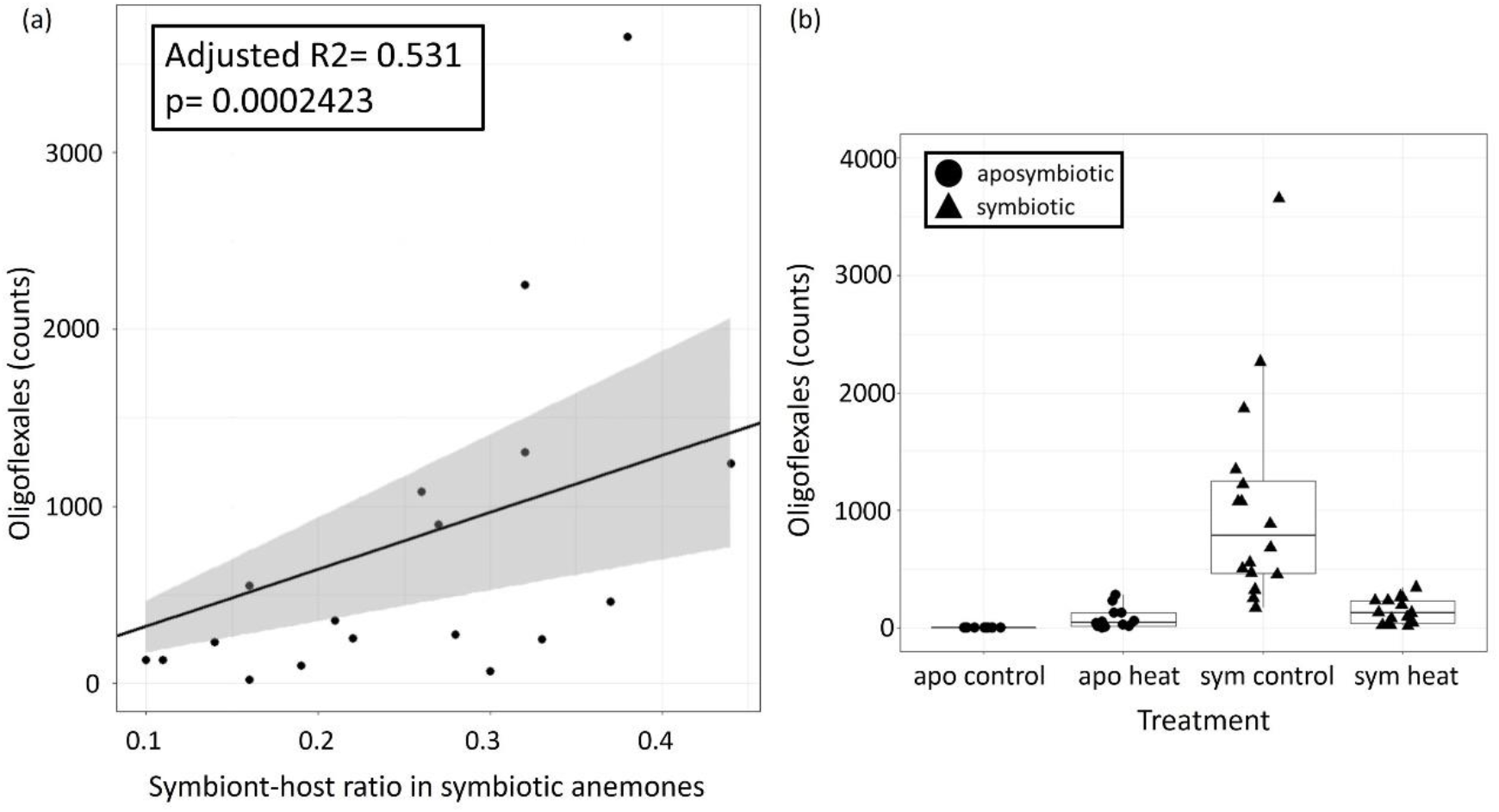
Oligoflexalcs in the microbiomc of Aiptasia. (a) Regression line through the origin and linear model results testing the relationship between the dependent variable, Oligoflexales bacterial counts (y axis) and explanatory variable, symbiont to host ratio (S/H) (x axis), (b) Total counts of Oligoflexales in rarefied data, by treatment group (black circles= aposymbiotic anemones, black triangles= symbiotic anemones).

### 3.3 Oligoflexales order are are associated with symbiotic state and are lost under thermal stress

Oligoflexales abundances were elevated in symbiotic anemones under control conditions but marginally elevated under heat treatment in aposymbiotic anemones relative to aposymbiotic controls (Fig. 3a, 3b). Total abundance counts clearly displayed this categorical difference as well (Fig. 4b). In addition to these categorical differences in Oligoflexales abundance by anemone symbiotic state and treatment (Fig. A1, 3, 4b), we also observed quantitative differences in abundance as a function of algal symbiont density. Symbiont to host (S/H) cell ratios of symbiotic anemones decreased under thermal stress indicating mild bleaching (p=0.047, Fig. A3, Table A7). Total S/H ratios for aposymbiotic anemones averaged around 0 but values slightly increased in anemones exposed to thermal stress (Fig. A3, Table A4).

A correlation matrix was built to examine pairwise relationships between the S/H ratio of symbiotic Aiptasia, alpha diversity, and Oligoflexales abundance counts. A moderate correlation between S/H ratio and the abundance of Oligoflexales bacteria was detected (Pearson’s Correlation, p=0.013, Fig. A4, Table A8). Regression analysis verified this positive correlation, revealing that 53% of the observed variation in Oligoflexales abundance across samples was explained by differences in S/H ratio (p < 0.001, Fig. 4a).

### 3.4 Oligoflexales may diversify under thermal stress

A phylogenetic tree plotting Oligoflexales ASV abundance showed symbiotic anemones in control conditions initially hosted 9 distinct ASVs but lost 1 after thermal stress (Fig. A5). In contrast, aposymbiotic anemones only harbored 2 ASVs under control conditions, whereas 8 Oligoflexales ASVs were detected in aposymbiotic anemones which experienced thermal stress (Fig. A5). We conducted a more detailed phylogenetic analysis to further investigate relationships among these distinct ASVs and other Oligoflexales variants identified in prior studies. Oligoflexales ASVs from this study grouped closely with three uncultured bacterium clones originating from an unrelated study on Aiptasia clonal strains CC7 and H2 (Randle *et al*., 2020) and one uncultured proteobacterium clone (GenBank: FJ425635) found in the microbiome of the scleractinian coral *Orbicella* (formerly *Montastrea*) *faveolata* (Fig. A6). Other Oligoflexales representatives were more distantly related (Fig. A6).

## DISCUSSION

Overall, understanding the ecological dynamics of algal-microbe interactions in cnidarians may be important for developing strategies to mitigate the impacts of climate change (Matthews *et al*., 2020). (Frommlet *et al*., 2015) found that a diverse community of bacteria facilitated the formation of symbiolites (spheroid, aragonite structures) in ex-hospite Symbiodiniaceae cultures. This discovery highlighted a unique approach that could potentially aid in coral calcification within reefs. Similarly, many eukaryotic algae are auxotrophs for the prokaryote-produced B vitamins, like B12, and must obtain B12 from symbionts or dietary sources for proper metabolic function and growth (Croft, Warren and Smith, 2006; Helliwell *et al*., 2011; Grossman, 2016). Furthermore, microbiome and algal symbiont co-occurrences may be a result of metabolic cooperation, as dinoflagellates may rely on necessary metabolites produced by bacteria and vice-versa (Cruz-López and Maske 2016; Grossman 2016; Kurihara et al. 2013). Additional possible functional roles of bacteria associated with Symbiodiniaceae may also span DMSP production, enhancing iron bioavailability and sulfur cycling (Lawson et al. 2018).

Here, we investigated the ecology of microbial communities, associated with Aiptasia, and how they are affected by presence of the algal symbiont and elevated temperatures. Despite treatment, Alphaproteobacteria and Gammaproteobacteria remained the predominant microbial taxa. Differences in beta diversity were observed between symbiotic and aposymbiotic animals under both control and thermal stress conditions but greater community similarity was observed among microbial populations in both symbiotic and aposymbiotic anemones exposed to heat stress. Additionally, elevated temperature decreased species richness in symbiotic anemones.We also show that Oligoflexales bacteria are part of the rare microbiome in symbiotic anemones but significantly decreased in abundance following thermal stress. The abundance of Oligoflexales was positively correlated with higher S/H cell ratio indicating symbiont density, rather than heat stress per se, impacted their abundance in Aiptasia.

### Exploring the cnidarian-algal-bacteria tripartite symbiosis, Oligoflexales as primary associates of Symbioidiniaceae

We examined the role of symbiotic algae in recruiting microbial taxa that may be specific to the microbiomes of symbiotic Aiptasia and identified two taxa (Oligoflexales and *Methylophilaceae*) that were significantly abundant in symbiotic Aiptasia only and decreased in relative abundance with symbiont loss both independent of and as a result of thermal stress (Fig. 3b,3d; 4). We focused on exploring Oligoflexales because these taxa were present in higher relative abundances and had been previously reported as associates of Aiptasia (Randle *et al*., 2020; Maire, Blackall and van Oppen, 2021). Whereas we could not find a consistent record of *Methylophilaceae* in symbiotic Aiptasia. Despite numerous surveys conducted on the Aiptasia microbiome (Röthig *et al*., 2016; Herrera *et al*., 2017; Ahmed *et al*., 2019; Hartman, van Oppen and Blackall, 2020; Costa *et al*., 2021), Oligoflexales remained undetected until recent research reported their presence in Aiptasia strain CC7 (Randle *et al*., 2020) and in the microbiome of Aiptasia acontia in strains AIMS1, AIMS2, AIMS3, and AIMS4 (Maire, Blackall and van Oppen, 2021). Two factors likely account for these results: (1) the utilization of identical sequencing primer sets across all three studies, which document the existence of Oligoflexales, including our own, and (2) the classification of taxonomy based on the most recent release of the SILVA database (v138, issued in 2019). We chose to use the 784F/1061R primer set because it captures global bacterial diversity while exhibiting low amplification of chloroplast and mitochondrial host DNA (Bayer *et al*., 2013; Andersson *et al*., 2008; Bayer *et al*., 2013) (Bayer *et al*., 2013). Furthermore, as Oligoflexales were recently recognized as a novel order under the Bdellovibrionota phylum (Nakai *et al*., 2014; Waite *et al*., 2020)), older databases may designate them as “unclassified/uncultured bacterial clones”. Based on our research and the prior studies, we conclude that Oligoflexales are a consistent component of the symbiotic microbiome in Aiptasia.

Bacteria belonging to the Oligoflexales order are Gram-negative, oligotrophic spirochaetes, and only one species, *Oligoflexus tunisiensis*, has been described and isolated (Nakai *et al*., 2014, 2016). We conducted a phylogenetic analysis of nine Oligoflexales ASV sequences against *O. tunisiensis* and other members of the parent phylum, Bdellovibrionota (Waite *et al*., 2020). Our ASVs formed a sister clade to *O. tunisiensis* and *Pseudobacteriovorax antillogorgiicola*, an isolate from gorgonian corals in the family Pseudobacteriovoracacea (McCauley, Haltli and Kerr, 2015) (Fig. A6). This suggests a close relationship between them. Currently, Pseudobacteriovoracacea are classified as Bdellovibrionales, but a proposal to reclassify them as Oligoflexales was submitted (Hahn *et al*., 2017), which is consistent with our findings grouping them with *O. tunisiensis*.

Although the ecological role of Oligoflexales in symbiotic Aiptasia remains a mystery, it has previously been suggested to comprise a set of taxa that aid thermotolerance in high salinities (Randle *et al*., 2020). The genome sequence of *O. tunisiensis* also provides clues on possible metabolic capabilities in Oligoflexales. Nakai et al. (2016) observed an incomplete denitrification pathway in *O. tunisiensis*, which resulted in the conversion of nitrate/nitrite (NO_3_/NO_2_) to nitrous oxide (N_2_O). Heterotrophic bacteria are known to recycle fixed nitrogen from the environment through denitrification, and those with a complete pathway can reduce fixed nitrogen to dinitrogen (N_2_) gas (Knowles, 1982). Nitrogen recycling by the host, Aiptasia, regulates algal symbiotic density (Cui *et al*., 2019) but bacterially regulated nitrogen may play a role in maintenance of the cnidarian-algal symbiosis as well, since nitrogen cycling is a hallmark of reciprocal bacterial association in cnidarian holobionts (Knowlton and Rohwer, 2003; Peixoto *et al*., 2017). It is unclear if the Oligoflexiales in the present study possess a similar denitrification pathway, but the genomic evidence from *O. tunisiensis* suggests that further research is needed to investigate this possibility.

We observed a higher abundance of Oligoflexales ASVs in symbiotic Aiptasia under control conditions, compared to symbiotic animals in the heat-stress treatment. In contrast, aposymbiotic Aiptasia in the control showed only 1 ASV but experienced an increase in diversity and abundance of ASVs in thermally stressed, aposymbiotic anemones (Fig. A5). While superficially this observation initially appears to contradict the notion that Oligoflexiales are associates of Symbiodiniaceae, we believe this pattern can be explained by the dynamics of symbiont maintenance and proliferation in Aiptasia (Jinkerson *et al*., 2022) and is actually fully consistent with the primary algal symbiont hypothesis.

Aposymbiotic Aiptasia anemones can retain remnant populations of algal symbionts, even in the absence of light. *S. linucheae* do not need to photosynthesize to be maintained in Aiptasia in dark conditions (Jinkerson *et al*., 2022). Jinkerson et al (2022) also found that *S. linuchae* did not proliferate *in-hospite* in the dark, but algal cell density significantly increased after subsequent transition to light conditions. We theorize that a small, non-detectable (by PCR) population of algal symbionts remained in the aposymbiotic group, despite continuous darkness for three months. This *S. linucheae* population likely proliferated during the combined light and heat-stress periods as confirmed by the slight increase in S/H ratios in this population (Fig. A3, Table A4), in contrast to the aposymbiotic controls only exposed to light. A recent study showed significantly higher proliferation of *S. linucheae* in Aiptasia at 32 °C, after 28 days compared to ambient temperatures (25 °C), suggesting cell division rates initially increased in response to elevated temperature but then declined after 12 weeks of sustained thermal stress (Herrera *et al*., 2021). The slight increase in algal abundance in the aposymbiotic population subjected to thermal stress was concomitant with an increase and diversification of Oligoflexales (Fig 4b, Fig A5). Whereas the opposite pattern (Oligoflexales loss) was observed in symbiotic Aiptasia exposed to heat stress. S/H ratios were lower (Table A4, Fig. A3) in symbiotic anemones exposed to thermal stress, and significant differences were observed between treatments (Table A7), indicating symbiont loss. Thermal stress can promote bleaching in symbiotic cnidarians leading to dysbiosis (Weis, 2008; Wooldridge, 2009)) but the mechanisms are not thoroughly established. One theory suggests dysbiosis may occur as a cascade effect that begins with a decrease in the photosynthetic efficiency of the symbiont, followed by a reduction in ammonium assimilation by the host, leading to an increase in the available ammonium pool. This, in turn, stimulates algal growth, and eventually, the host becomes unable to keep up with the resulting internal metabolic shifts, leading to the expulsion of the symbionts (Cui *et al*., 2019; Rädecker *et al*., 2021). We hypothesize that heat-stressed, aposymbiotic Aiptasia did not experience the same metabolic constraints and were able to adequately support growth of algal populations, leading to slightly higher symbiont abundance and increased abundance/diversity of Oligoflexales bacteria (Fig. 4b, Fig. A5).

Lastly, it is a possibility that Oligoflexales observed in aposymbiotic anemones under heat conditions were environmentally acquired. We used 0.2 µm FSW to maintain our cultures but previous research has demonstrated that *O. tunisiensis* is part of the 0.2 µm filtrate culturable fraction (Nakai *et al*., 2014). Although *O. tunisiensis* can reach up to 10 µm in length, and are between 0.4 - 0.8 µm wide, they can compress through 0.2 µm filter pores. We do not have morphological data for the Oligoflexales in this study, so we cannot eliminate the possibility of environmental contamination. However, all animals were clonal propagates and were maintained using 0.2 µm FSW, originating from the same 20 L tank, yet displayed distinct microbiome profiles according to treatment (Fig 3). Furthermore, the disparity of Oligoflexales abundances between treatment groups (Fig. 4b) indicates symbiotic-state specificity in control conditions.

Environmentally acquired or not, Oligoflexales may play an important role in the holobiont as endosymbiotic partners that only associate with cnidarians when microalgal symbionts are present, as rare members of the symbiotic microbiome. It is possible that Oligoflexales, like rare members of the coral core microbiome, are widespread in anemone tissue, particularly in close proximity to algal symbionts (D Ainsworth *et al*., 2015) and future work should aim to fully characterize their abundance and distribution.

### Microbiome assemblages of aposymbiotic and symbiotic Aiptasia

Microbial assemblages in Aiptasia from wild and cultured populations (including clonal strain CC7) in ambient conditions showed consistent similarity at the phylum level but not at lower taxonomic levels (Brown *et al*., 2017). In all our samples, regardless of treatment, *Alcanivorax* sp. from the Oceanospirillales order, *Cognatishimia* sp., and an unclassified bacterium from the Rhodobacteraceae family from the Rhodobacterales order accounted for around 60-85% of the total relative abundance (Fig. A2). This observation, that 2-3 taxa are numerically dominant in Aiptasia, contradicts other microbial surveys on Aiptasia that identified a wider range of taxonomic groups within the same relative abundance range (Röthig *et al*., 2016; Randle *et al*., 2020; Curtis *et al*., 2023). The importance of functional redundancy in shaping microbial communities in Aiptasia is emphasized by the conflicting results obtained from various studies. Functional redundancy refers to a diverse range of bacteria with similar capabilities, able to perform similar functions in the same niche (Louca *et al*., 2018). Functional redundancy is an advantageous strategy for ecosystem stability and may play a role in the resilience of hosts, like corals, facing environmental disturbances (Voolstra and Ziegler, 2020).

Prior work conducted by (Röthig *et al*., 2016; Ahmed *et al*., 2019; Randle *et al*., 2020) showed that aposymbiotic and symbiotic Aiptasia (CC7), hosted distinct microbiomes. Here we expanded upon this finding by introducing a stressor to assess the response of the aposymbiotic microbiome and the symbiotic microbiome under thermal stress (Fig. A1). Although alpha-diversity did not differ between symbiotic status (Fig. 2a), beta-diversity was dissimilar (Fig. 2b, Table A6), which prompted us to explore which taxa were responsible for these observations.

We observed a significant difference in the relative abundance of Oligoflexales, Saccharospirillaceae, Pseudoalteromonadaceae, and Methylophilaceae families between symbiotic and aposymbiotic animals, with these taxa being more prevalent in the former (Fig. 3b). Our findings align with other studies that have demonstrated differences in relative abundance of microbiome composition between aposymbiotic and symbiotic states in strain CC7 (Röthig *et al*., 2016; Sydnor, 2020; Curtis *et al*., 2023). Nevertheless, the four differentially abundant taxa we identified were not previously reported, suggesting genotype by environment and symbiotic state all influence microbial assemblages in Aiptasia.

### Microbiome fluctuations caused by heat stress changed the rare microbiome

Here, we aimed to identify microbial taxa that are linked with symbiotic states in a defined clonal strain of Aiptasia and tracked microbial fluctuations of these taxa following mild bleaching. The dominant taxa in all Aiptasia were found to be unaffected by both temperature and symbiotic state. However, the rare microbiome of symbiotic Aiptasia was significantly affected by temperature stress. In contrast to the other treatment groups, symbiotic Aiptasia exhibited a reduction in alpha-diversity and beta-diversity convergence after six days of exposure to heat stress (Fig. 2). A decrease in alpha-diversity was also observed in another study exposing symbiotic Aiptasia CC7 to short-term heat stress (Sydnor, 2020), but it appears to be a temporary phenomenon since a long-term study by (Ahmed *et al*., 2019)) on Aiptasia CC7, surveying microbial communities under continuous heat stress (32°C for two years) demonstrated an increase in both alpha-diversity and number of bacterial taxa relative to paired controls. Here, symbiotic Aiptasia experienced microbiome restructuring when subjected to heat stress (Fig. 3d) and the resulting assemblage was most similar to that observed in heat-stressed aposymbiotic anemones (Fig. 3c). This suggests that short-term heat stress in Aiptasia may be the main driver that led to a convergence of microbial communities in both aposymbiotic and symbiotic animals.

However, possible symbiont gains in the aposymbiotic and symbiont loss in the symbiotic anemones exposed to thermal stress (Fig. A3) may also be contributing to the convergence of microbial communities. Symbiotic anemones exhibited increased relative abundance of several bacteria taxa, most noticeably Oligoflexales, Saccharospirillaceae, and Pseudoalteromonadaceae compared to aposymbiotic anemones under control temperatures (Fig. 3b). These same taxa increase in relative abundance in heat-stressed aposymbiotic anemones compared to control aposymbiotic anemones (Fig. 3a) until their relative abundance is indistinguishable from those observed in the symbiotic anemones in response to thermal stress (Fig. 3c). Whereas their abundances decrease in response to heat stress in symbiotic anemones (Fig. 3d). The most parsimonious explanation for these apparently contradictory changes in response to heat stress is that it is not thermal stress, but algal symbiont density which influences patterns of convergence. Additional time-course studies examining repopulation of algal symbiont communities would provide additional support for this hypothesis.

It is unclear whether changes in the relative abundance or composition of the microbial community affected the physiology or fitness of aposymbiotic and symbiotic Aiptasia hosts as we did not conduct any host-specific assays, but we can confirm that no Aiptasia died during the course of this experiment. While we have identified a strong association between the abundance of Oligoflexales and algal endosymbionts independent of thermal stress, whether these bacteria are mutualists or commensals remains unresolved. Additional work on patterns of localization, metabolic exchange, and spatial and temporal fidelity are needed. But given the tractability of the Aiptasia model, this represents a promising future study system for investigating multi-partner symbioses. Understanding how the absence of a member in a multipartite symbiosis impacts the resilience of other organisms in the holobiont can uncover valuable insights into how symbiotic organisms respond to environmental challenges.

## ACKNOWLEDGEMENTS

This study was funded by the National Science Foundation Graduate Research Fellowship Program grant award DGE-1418060 to EGA and start-up funding from the University of Southern California to CDK. We would like to thank Dr. Cory J Krediet for providing a set of clonal Aiptasia CC7 anemones. Special thanks to Dr. Ross Cunning, Dr. Sheila Kitchen and Dr. Angela Poole for invaluable assistance in determining appropriate qPCR primers. Additionally, we would like to thank Daniel Olivares-Zambrano for help pooling the libraries before sequencing and Maria Ruggeri for kindly providing Aiptasia CC7 DNA samples to test the qPCR primers used to calculate symbiont to host ratios.

## CONFLICT OF INTEREST

The authors declare no competing interests.

## AUTHOR CONTRIBUTIONS

EGA and CDK conceived and designed the field experiment and obtained funding. EGA and MJF performed anemone maintenance, aposymbiotic generation and sampling for the experiment. EGA completed DNA extractions, library preparation, qPCR and all bioinformatic and statistical analyses and wrote the first draft of the manuscript. All authors contributed to revisions.

## DATA ACCESSIBILITY

Scripts for data analysis and statistical output for differential heat trees generated using the Metacoder R package are archived at Zenodo, <u>doi.org/10.5281/zenodo.7693398</u>. It can also be viewed at https://github.com/symbiotic-em/aiptasia_thermal_stress. Demultiplexed sequences are available at the National Center for Biotechnology Information (NCBI) Sequence Read Archives (SRA) under accession code: <u>PRJNA929535</u>.

## Notes

https://github.com/symbiotic-em/aiptasia_thermal_stress

https://zenodo.org/record/7693398#.ZDnnwLrMJPY

https://www.ncbi.nlm.nih.gov/bioproject/PRJNA929535

### Competing Interest Statement

The authors have declared no competing interest.

