## appendix for "Abundance of Oligoflexales bacteria is associated with algal symbiont density independent of thermal stress in Aiptasia anemones"

Figures A1-A6

Tables A1-Table A8

#### ABSTRACT

Many multicellular organisms, such as humans, plants, and invertebrates, depend on symbioses with microbes for metabolic cooperation and exchange. Reef-building corals, an ecologically important order of invertebrates, are particularly vulnerable to environmental stress in part because of their nutritive symbiosis with dinoflagellate algae, and yet also benefit from these and other microbial associations. While coral microbiomes remain difficult to study because of their complexity, the anemone Aiptasia is emerging as a simplified model. Research has demonstrated co-occurrences between microbiome composition and the abundance and type of algal symbionts in cnidarians. However, whether these patterns are the result of general stress-induced shifts or depletions of algal-associated bacteria remains unclear. Our study aimed to distinguish the effect of changes in symbiont density and thermal stress on the microbiome of symbiotic Aiptasia strain CC7 by comparing them with aposymbiotic anemones, depleted of their native symbiont, *Symbiodinium linucheae*. Our analysis indicated that overall, thermal stress had the greatest impact on disrupting the microbiome. We found that three bacterial classes made up most of the relative abundance (60-85 %) in all samples, but the rare microbiome fluctuated between symbiotic states and following thermal stress. We also observed that *S. linucheae* density correlated with abundance of Oligoflexales, suggesting these bacteria may be primary symbionts of the dinoflagellate algae. The findings of this study help expand knowledge on prospective multipartite symbioses in the cnidarian holobiont and how they respond to environmental disturbance**.**

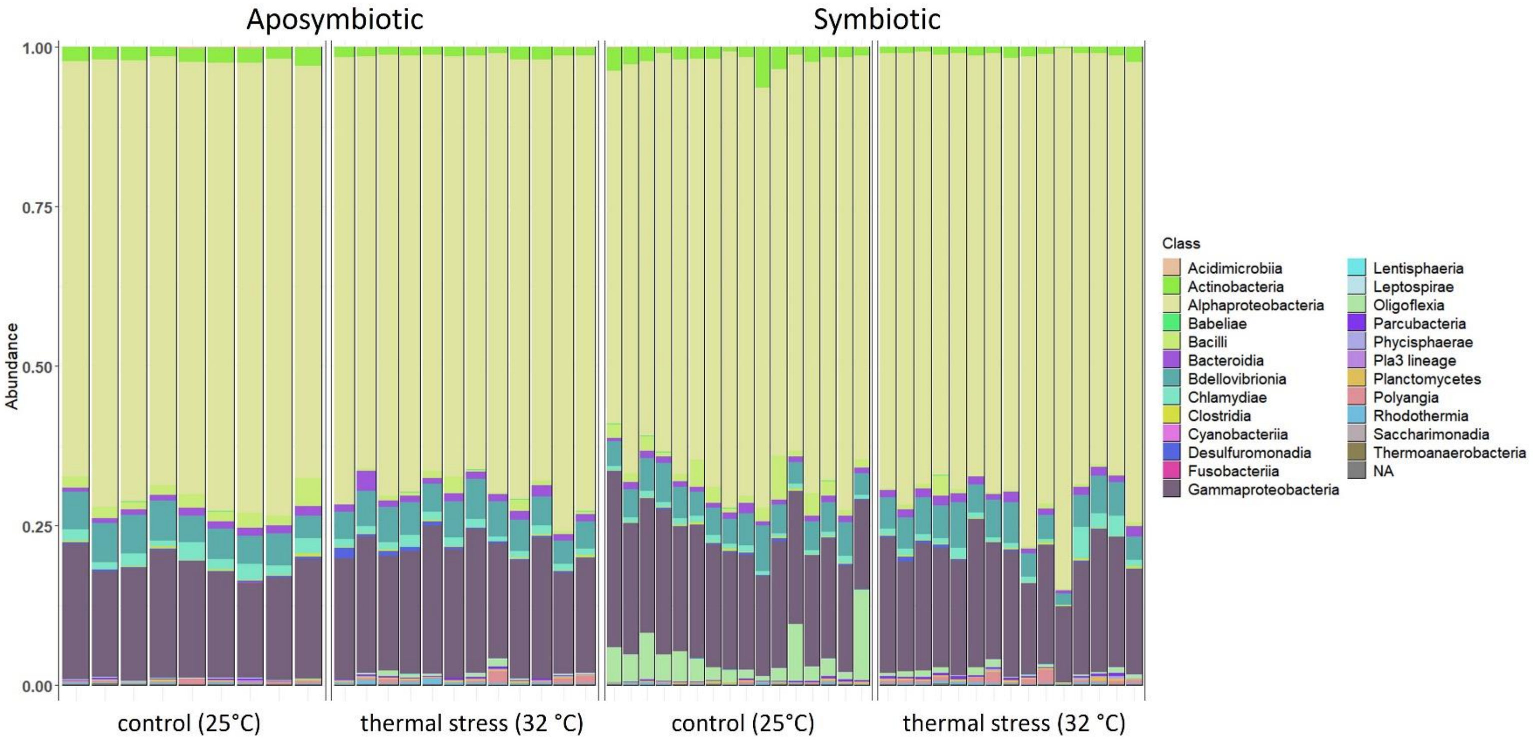


**Figure A1**. Relative abundance barplots of taxa, by class, in aposymbiotic and symbiotic anemones by experimental conditions (25 °C vs 32 °C). Alphaproteobacteria, Bdellovibrionota and Gammaproteobacteria dominate all the samples. Oligoflexia are distinctly present in control symbiotic anemones.

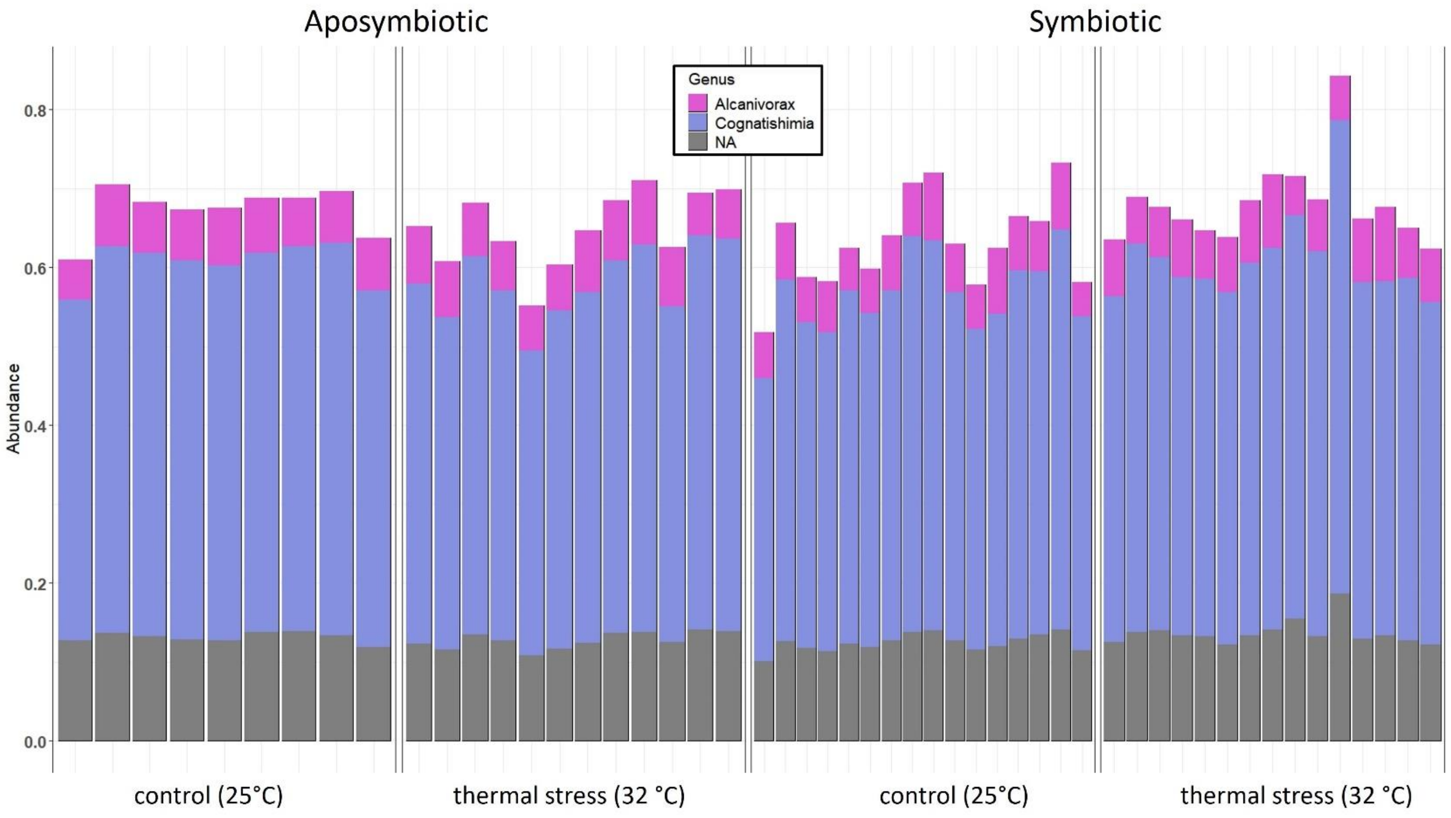


**Figure A2.** Relative abundance of the three most abundant genera in all *Aiptasia* groups: *Alcanivorax* belonging to the Oceanospirillales order, *Cognatishimia* and an unclassified bacterium, both genera are part of the Rhodobacterales order.

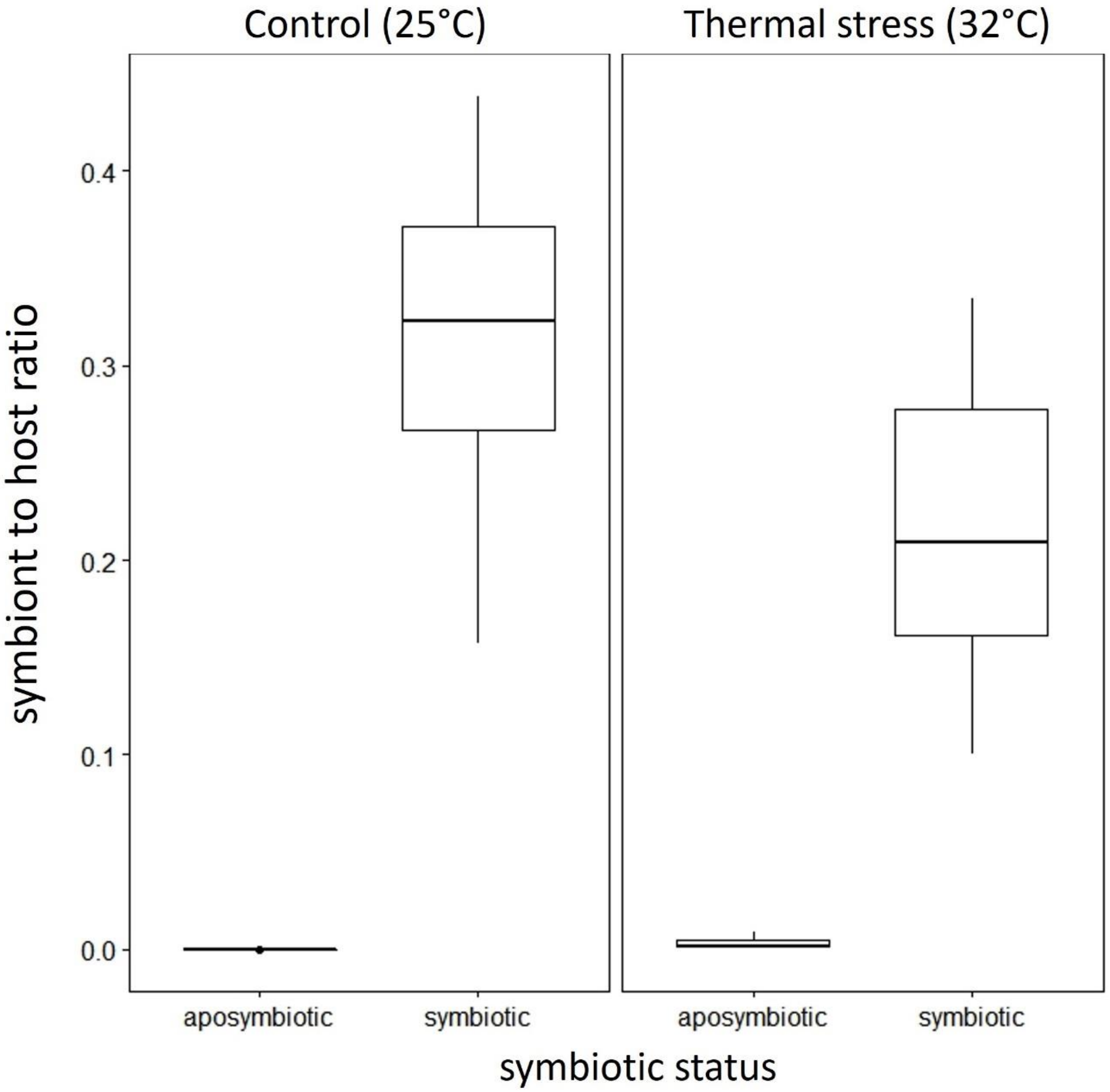


**Figure A3.** Symbiont to host ratio (S/H) in *Aiptasia* anemones. Ribosomal protein L10 was used as a reference for *Aiptasia* and actin locus gene for *Symbiodinium*. In both treatments, aposymbiotic *Aiptasia* approximated zero but ratios in symbiotic *Aiptasia* decreased after mild thermal stress (p= 0.047).

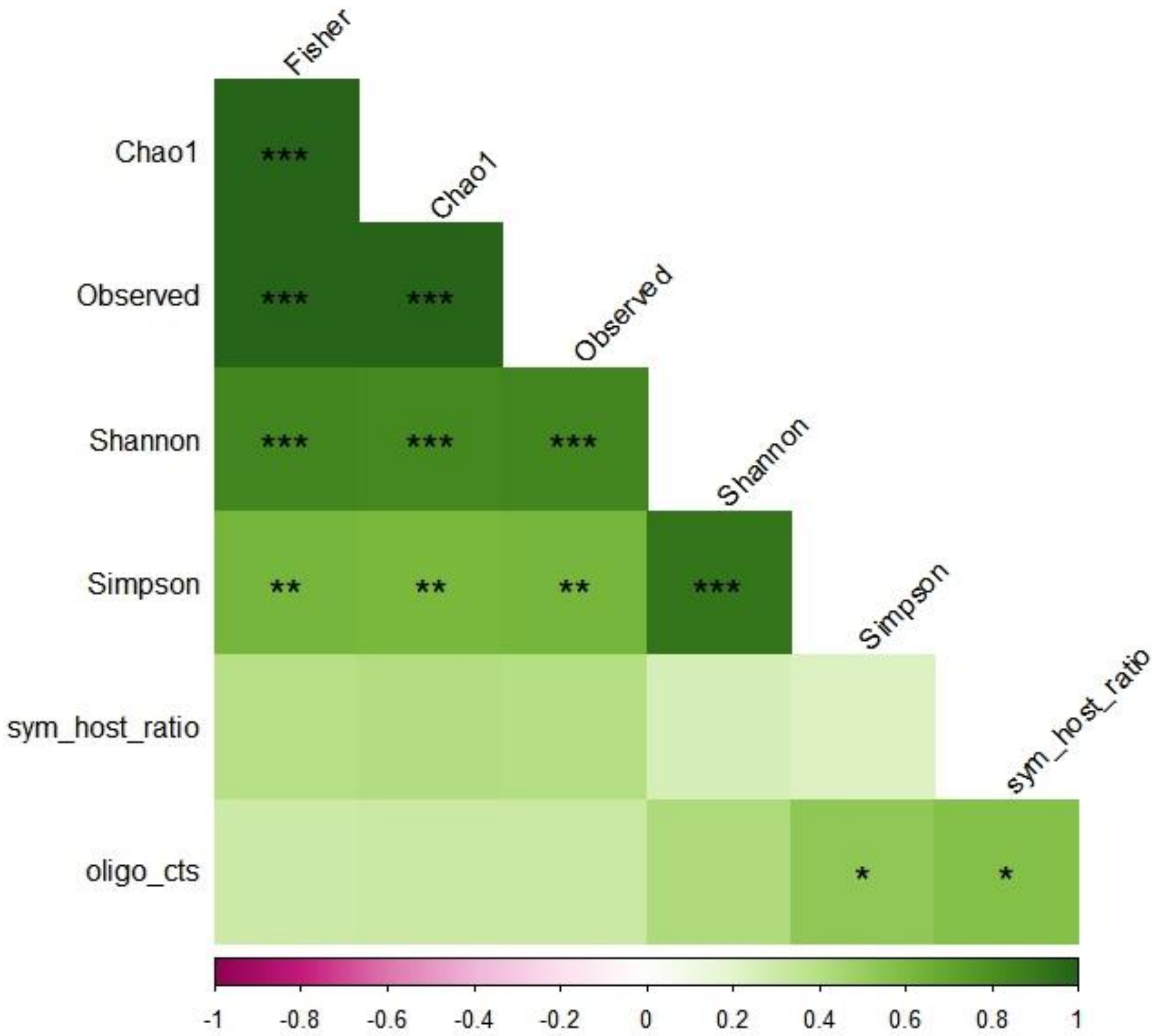


**Figure A4.** Correlation matrix showing the strength of interactions between S/H ratio ("sym_host_ratio") and a set of variables containing six alpha diversity indices (Fisher, Chao1, Shannon, Simpson) and Oligoflexales (oligo_cts) counts. S/H ratio displays a significant correlation with one variable, Oligoflexales abundance.

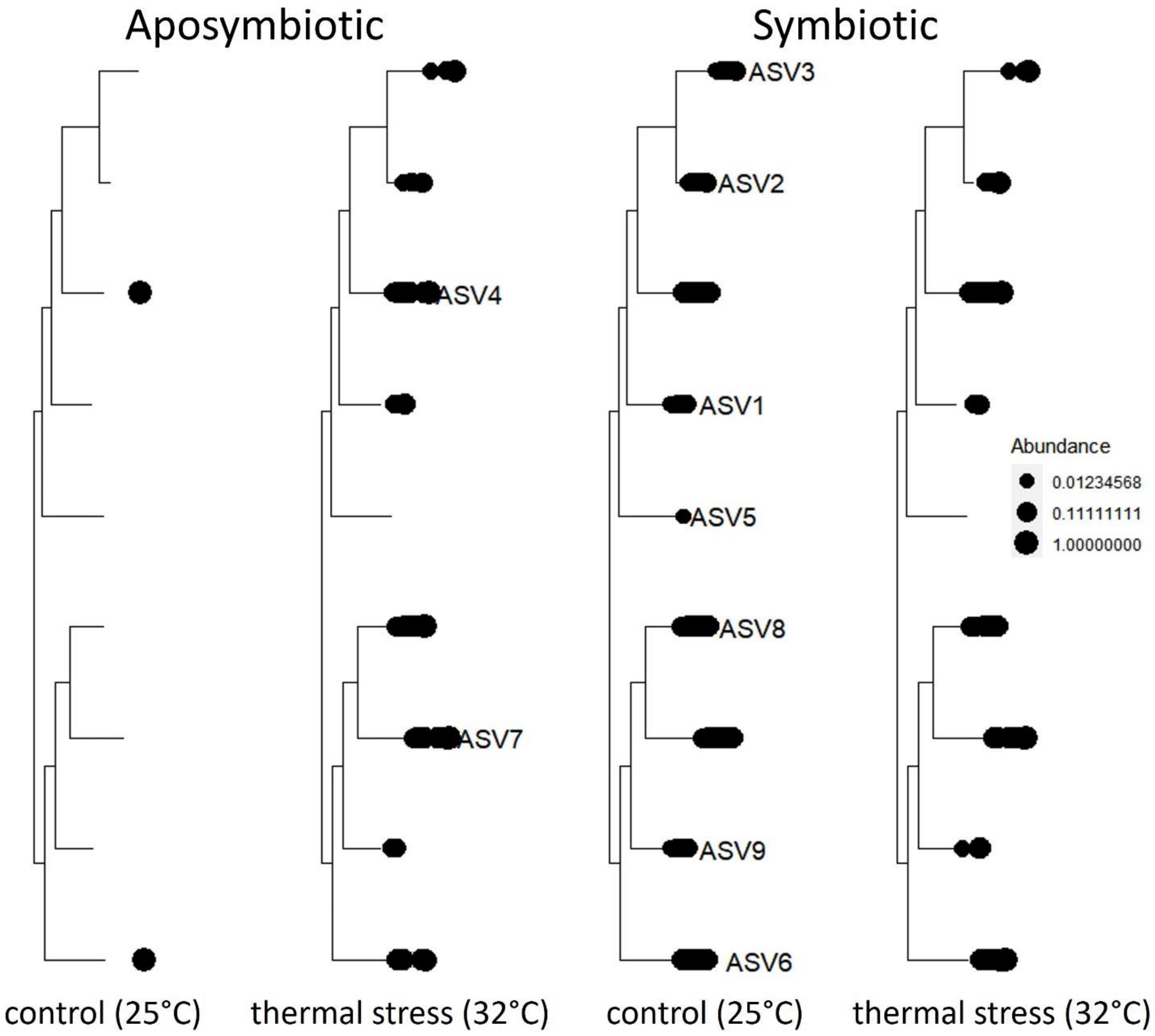


**Figure A5.** Phylogenetic visualization of Oligoflexales ASV abundance per *Aiptasia* treatment group (aposymbiotic control, aposymbiotic thermal stress, symbiotic control, and symbiotic thermal stress) using rarefied and filtered data. A total of 9 ASVs were observed. ASV 4 and ASV 6 were observed in all samples but differed in abundance. ASV abundance and diversification in aposymbiotic anemones increased in heat stressed anemones but decreased in symbiotic anemones.

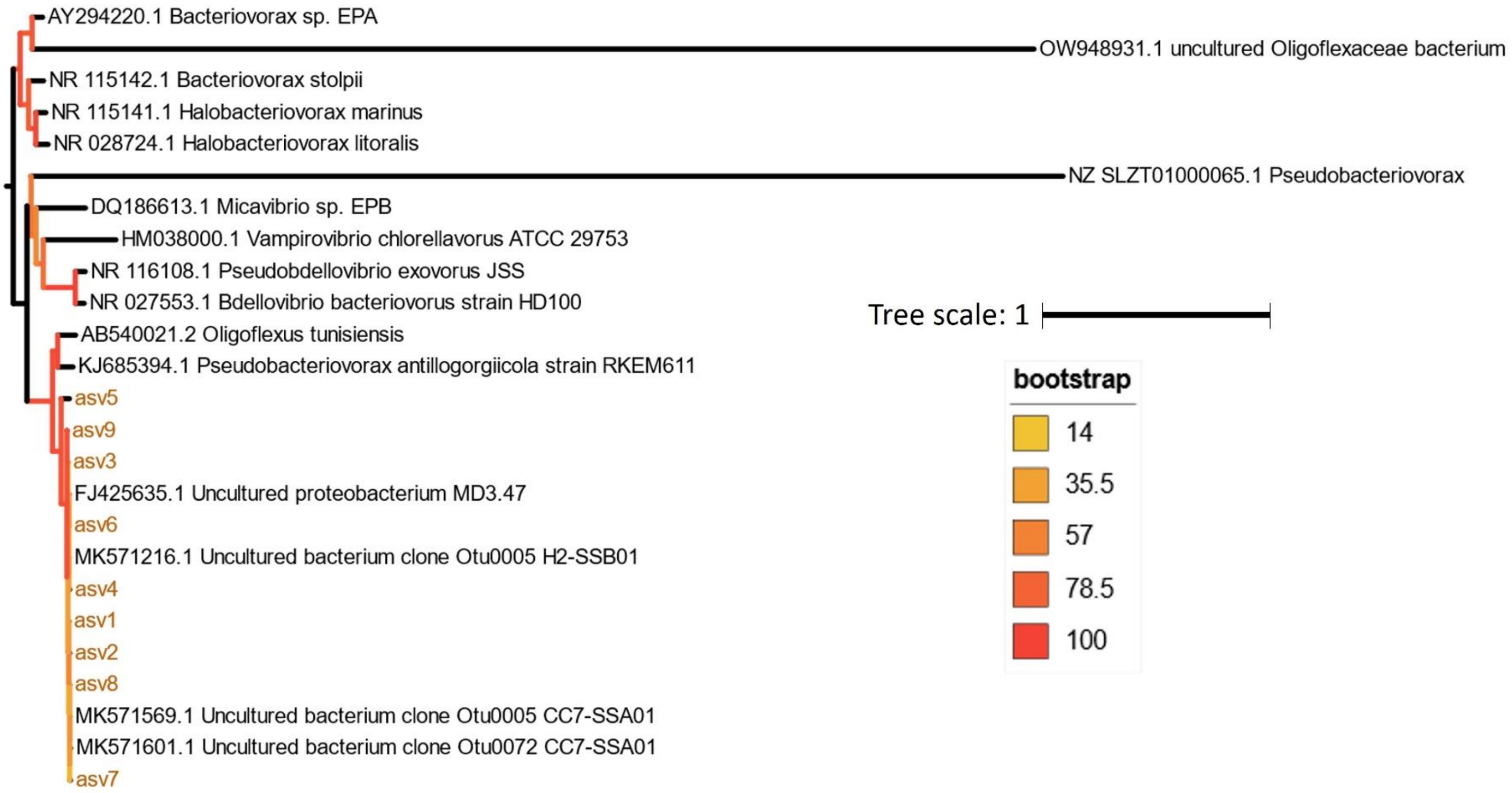


**Figure A6.** Oligoflexales phylogenetic analysis. Maximum likelihood phylogeny with ultrafast bootstrap (n=1,000 replicates) of this study's (golden color) Oligoflexales ASVs with two published Oligoflexales sequences (AB540021.2 and OW948931.1), along with four sequences that were close matches to our ASV sequences (FJ425635.1, MK571216.1, MK571569.1, MK571601.1) and members of the Bdellovibrionota phylum (*Bdellovibrionia* and *Bacterovoracia*). Surprisingly, an uncultured Oligoflexaceae bacterium (OW948931.1) did not cluster with the other confirmed member of Oligoflexiales (AB540021.2, *Oligoflexus tunisiensis*) or presumptive members of Oligoflexiales (ASVs in this study) but clustered with a *Bacteriovorax* sp.

**Table A1**. Read/ASV pipeline using DADA2 and Phyloseq.

DADA2/ Phyloseq ASV Pipeline

| 52 samples | DADA2 sequence table | chloroplast and mitochondria removal (Phyloseq) | rarefaction at 26, 481 reads | Filtered taxa (occur 3x in more than 4 samples) |
|---|---|---|---|---|
| Reads | 4,996,356 | 4,984,116 | 1,377,012 | 1,338,082 |
| ASVs | 4,577 | 4,471 | 4,299 | 774 |

**Table A2.** Significant, consistent matches to our 9 queried Oligoflexales ASVs, using standard NCBI BLAST (blastn suite). GenBank MK571601.1, MK571569.1 and MK571216.1 originate from (Randle et al. 2020). GenBank FJ425635 derives from an unpublished study on the microbiome of the coral *Orbicella* (formerly *Montastrea) faveolata*.

| **Query** | **MK571601.1** | | **MK571569.1** | | **MK571216.1** | | **FJ425635** | |
|---|---|---|---|---|---|---|---|---|
| **Oligoflexales ASV** | Percent identity (%) / bit score | e-value | Percent identity (%) / bit score | e-value | Percent identity (%) / bit score | e-value | Percent identity (%) / bit score | e-value |
| 1 | 99.23/ 468 | 3.00E-127 | 99.61/ 473 | 6.00E-129 | 99.61/ 473 | 6.00E-129 | 99.66/ 534 | 3.E-147 |
| 2 | 99.23/ 468 | 3.00E-127 | 99.61/ 473 | 6.00E-129 | 99.61/ 473 | 6.00E-129 | 99.66/ 534 | 3.E-147 |
| 3 | 99.23/ 468 | 3.00E-127 | 99.61/ 473 | 6.00E-129 | 99.61/ 473 | 6.00E-129 | 99.66/ 540 | 6.E-149 |
| 4 | 99.61/ 473 | 6.00E-129 | 100/ 479 | 1.00E-130 | 100/ 479 | 1.00E-130 | 100/ 542 | 2.E-149 |
| 5 | 94.62/ 401 | 3.00E-107 | 95/ 407 | 6.00E-109 | 95/ 407 | 6.00E-109 | 95.58/ 470 | 8.E-128 |
| 6 | 99.61/ 473 | 6.00E-129 | 100/ 479 | 1.00E-130 | 100/ 479 | 1.00E-130 | 100/ 545 | 1.E-150 |
| 7 | 99.61/ 473 | 6.00E-129 | 100/ 479 | 1.00E-130 | 100/ 479 | 1.00E-130 | 100/ 540 | 6.E-149 |
| 8 | 99.61/ 473 | 6.00E-129 | 100/ 479 | 1.00E-130 | 100/ 479 | 1.00E-130 | 100/ 540 | 6.E-149 |
| 9 | 99.23/ 468 | 3.00E-127 | 99.61/ 473 | 6.00E-129 | 99.61/ 473 | 6.00E-129 | 99.66/ 536 | 7.E-148 |

**Table A3.** Samples with sufficient DNA for qPCR analysis to determine S/H ratio. After qPCR analysis, three "symbiotic" samples were omitted from downstream analysis due to incompatibility with the 16S rRNA dataset (these three samples produced low Illumina read yields and were filtered from the sequencing dataset).

Symbiont to Host (S/H) Ratio Sample Pipeline

| | Initial samples | Samples with enough DNA for qPCR | Samples used to determine S/H ratio | Samples used for statistical analysis |
|---|---|---|---|---|
| Aposymbiotic | 24 | 15 | 8 control, 7 heat | 0 |
| Symbiotic | 36 | 21 | 8 control, 10 heat | 8 control , 10 heat |

**Table A4**. Average cycle threshold (Ct) values of samples used for qPCR analysis. Primers L10 (host) and actin (symbiont) were used to calculate S/H ratio using the formula: $(2^{(Ct\ host/Ct\ sym)}) * 2$

| sample_ID | Primer-L10_ct | status | Primer-actin_ct | treatment | sym_host_ratio $(2^{\left(\frac{host\ ct}{sym\ ct}\right)}) * 2$ |
|---|---|---|---|---|---|
| c1-1 | 16.295 | aposymbiotic | 40 | control | 1.46E-07 |
| c1-2 | 16.95 | symbiotic | 19.58 | control | 0.323088 |
| c1-3 | 16.885 | symbiotic | 19.85 | control | 0.256139 |
| c1-4 | 15.985 | aposymbiotic | 40 | control | 1.18E-07 |
| c1-5 | 15.63 | aposymbiotic | 40 | control | 9.22E-08 |
| c1-6 | 16.82 | symbiotic | 19.01 | control | 0.438303 |
| c2-1 | 15.88 | symbiotic | 18.77 | control | 0.269807 |
| c2-1 | 16.305 | aposymbiotic | 40 | control | 1.47E-07 |
| c2-2 | 16.11 | aposymbiotic | 40 | control | 1.29E-07 |
| c2-3 | 16.145 | symbiotic | 18.585 | control | 0.368567 |
| c2-3 | 16.425 | aposymbiotic | 40 | control | 1.6E-07 |
| c2-4 | 16.045 | aposymbiotic | 40 | control | 1.23E-07 |
| c2-5 | 18.12 | aposymbiotic | 40 | control | 5.18E-07 |
| c3-1 | 15.41 | symbiotic | 18.045 | control | 0.32197 |
| c3-2 | 15.85 | symbiotic | 19.52 | control | 0.157127 |
| c3-5 | 16.98 | symbiotic | 19.375 | control | 0.380245 |
| blank | 29.1225 | blank | 31.34 | control | |
| 1-1 | 16.335 | symbiotic | 19.97 | thermal_32C | 0.160985 |
| 1-1 | 16.745 | aposymbiotic | 26.91 | thermal_32C | 0.001742 |
| 1-2 | 15.69 | symbiotic | 19.89 | thermal_32C | 0.108819 |
| 1-3 | 15 | symbiotic | 18.84 | thermal_32C | 0.139661 |
| 1-4 | 15.81 | symbiotic | 18.98 | thermal_32C | 0.222211 |
| 1-5 | 16.72 | symbiotic | 19.3 | thermal_32C | 0.334482 |
| 1-5 | 15.65 | aposymbiotic | 23.66 | thermal_32C | 0.007759 |
| 1-6 | 16.18 | aposymbiotic | 27.24 | thermal_32C | 0.000937 |
| 1-6 | 18 | symbiotic | 21.61 | thermal_32C | 0.163799 |
| 2-1 | 15.63 | symbiotic | 18.89 | thermal_32C | 0.208772 |
| 2-2 | 15.91 | aposymbiotic | 26.84 | thermal_32C | 0.001025 |
| 2-2 | 15.68 | symbiotic | 18.53 | thermal_32C | 0.277392 |
| 2-3 | 15.97 | aposymbiotic | 26.48 | thermal_32C | 0.001372 |
| 2-4 | 16.17 | symbiotic | 19.36 | thermal_32C | 0.219151 |
| 2-5 | 15.02 | aposymbiotic | 22.79 | thermal_32C | 0.009163 |
| 2-6 | 16.27 | symbiotic | 19.7 | thermal_32C | 0.185565 |
| 2-6 | 16 | aposymbiotic | 26.67 | thermal_32C | 0.001228 |
| 3-2 | 16.56 | symbiotic | 19.31 | thermal_32C | 0.297302 |
| 3-5 | 15.5 | symbiotic | 19.82 | thermal_32C | 0.100134 |
| 3-6 | 16.67 | symbiotic | 19.36 | thermal_32C | 0.309927 |
| blank | 36.8 | blank | 27 | thermal_32C | |

**Table A5.** Alpha diversity statistical output. A one-way ANOVA test was used to determine alpha diversity (Chao1 index) differences between groups. Post hoc analysis was performed, using Tukey multiple comparison of means (95% family-wise confidence level), to assess which groups are different from the rest.

ANOVA, p.value=0.009
Tukey multiple comparison of means
95% family-wise confidence level
Fit: aov(formula = alph$Chao1 ~ my_factors$grupos)

| | diff | lwr | upr | p. adj |
|---|---|---|---|---|
| apo_heat-apo_control | -16.754810 | -178.0198 | 144.5102193 | **0.9925194** |
| sym_control-apo_control | -5.001352 | -157.3825 | 147.3797773 | **0.9997576** |
| sym_heat-apo_heat | -141.616617 | -283.2573 | 0.0240811 | **0.0500538** |
| sym_heat-sym_control | -153.370075 | -284.8071 | -21.9330695 | **0.0162628** |

**Table A6.** Beta diversity statistical output using either the nonparametric test, PERMANOVA for assessing differences between *Aiptasia* microbial assemblages (symbiotic control vs symbiotic heat stressed, aposymbiotic control vs aposymbiotic heat stressed) with even group dispersions or ANOSIM for microbial assemblages with uneven dispersion (symbiotic control vs aposymbiotic control).

**PERMANOVA (adonis2 function)**

### of permutations: 999

- sym_ctrl_vs_sym_heat

| | Df | R2 | Pr(>F) |
|---|---|---|---|
| grupos | 1 | 0.19 | 0.001 *** |
| Residual | 29 | 0.47 | |

- apo_ctrl_vs_apo_heat

| | Df | R2 | Pr(>F) |
|---|---|---|---|
| grupos | 1 | 0.22 | 0.001 *** |
| Residual | 19 | 0.78 | |

**ANOSIM (anosim function)**

### of permutations: 999

- sym_ctrl_vs_apo_ctrl

ANOSIM Statistic R: 0.28
Significance: 0.003

**Table A7.** Statistical output of linear mixed effects model, fit by residual maximum likelihood (REML), to account for random effects (from tank differences between treatment groups).

**Treatment effect on S/H ratio, after accounting for tank as a random effect**

```
> summary(model1)
Linear mixed-effects model fit by REML
  Data: sym_df
       AIC       BIC logLik
  -22.03001 -18.93965 15.015

Random effects:
 Formula: ~1 | tank
        (Intercept)   Residual
StdDev: 1.783909e-06 0.08255301

Fixed effects:  sym_host_ratio ~ treatment
                    Value  Std.Error DF   t-value p-value
(Intercept)         0.315 0.02918690 12 10.792514  0.0000
treatmentthermal_32C -0.111 0.03915833  4 -2.834646  0.0471
 Correlation:
                    (Intr)
treatmentthermal_32C -0.745

Standardized Within-Group Residuals:
       Min          Q1         Med          Q3         Max
-1.87758137 -0.63595498  0.06056714  0.75708926  1.52629196

Number of Observations: 18
Number of Groups: 6
> anova(model1)
            numDF denDF   F-value p-value
(Intercept)     1    12 169.50844  <.0001
treatment       1     4   8.03522  0.0471
```

**Table A8.** Pearson's correlation p-values, corresponding to the correlation matrix on Fig. A6.

| Factor | p.sym_host_ratio | p.Chao1 | p.Observed | p.Shannon | p.Simpson | p.Fisher | p.oligo_cts |
|---|---|---|---|---|---|---|---|
| **sym_host_ratio** | 0 | 0.088541 | 0.09166 | 0.28477 | 0.34754 | 0.100708 | **0.013438** |
| Chao1 | 0.088541 | 0 | 9.20E-32 | 1.33E-05 | 0.00651 | 6.41E-25 | 0.208603 |
| Observed | 0.09166 | 9.20E-32 | 0 | 1.05E-05 | 0.00595 | 3.24E-26 | 0.207116 |
| Shannon | 0.28477 | 1.33E-05 | 1.05E-05 | 0 | 4.93E-08 | 9.44E-06 | 0.081825 |
| Simpson | 0.34754 | 0.00651 | 0.00595 | 4.93E-08 | 0 | 0.005939 | 0.023221 |
| Fisher | 0.100708 | 6.41E-25 | 3.24E-26 | 9.44E-06 | 0.005939 | 0 | 0.219931 |
| oligo_cts | 0.013438 | 0.208603 | 0.207116 | 0.081825 | 0.023221 | 0.219931 | 0 |
